# Sex-determination and sex chromosomes are shared across the radiation of dioecious *Nepenthes* pitcher plants

**DOI:** 10.1101/240259

**Authors:** Mathias Scharmann, T. Ulmar Grafe, Faizah Metali, Alex Widmer

## Abstract

Plants with separate sexes (dioecy) represent a minority but dioecy has evolved multiple times independently in plants. Our understanding of sex determination systems in plants and of the ecological factors and molecular changes associated with the evolution of dioecy remain limited. Here, we study the sex-determination system in dioecious plants that lack heteromorphic sex chromosomes and are not amenable to controlled breeding: *Nepenthes* pitcher plants. We genotyped wild populations of flowering males and females of three *Nepenthes* taxa using ddRAD-seq, and sequenced a male inflorescence transcriptome. We developed a novel statistical tool (privacy rarefaction) to distinguish true sex-specificity from stochastic noise in high-throughput sequencing data. Our results support XY-systems in all three *Nepenthes* taxa and in *Silene latifolia* which was used as a positive control for its known XY-system. The male-specific region of the Y chromosome showed little conservation among the three *Nepenthes* taxa, except for the essential pollen development gene DYT1 which was also male-specific in additional taxa. Hence, this homomorphic XY sex-determination system likely has a unique origin older than the crown of the genus *Nepenthes* at c. 17.7 My. In addition to the characterisation of the previously unknown sex chromosomes of *Nepenthes*, our work contributes an innovative, highly sensitive statistical method to efficiently detect sex-specific genomic regions in wild populations in general.

## Introduction

### Dioecy and sex chromosomes

Although the majority of flowering plant species have hermaphroditic flowers, plant sexual systems and mechanisms of sex-determination are highly diverse (Charlesworth 2002; Bachtrog et al. 2014). Only 5–6% of species have female and male flowers on separate individuals (dioecy), while the evolutionary transition to dioecy occurred more than 800 times independently in angiosperms alone (Renner 2014). In contrast to the outcrossing–selfing transition, for which many of the underlying genetic changes have recently been uncovered (Shimizu & Tsuchimatsu 2015), relatively little is known about the transitions from hermaphroditism to dioecy and the mechanisms of sex-determination in plants (Charlesworth 2016). The main hypotheses for the evolution of separate sexes in plants involve conflicting trait optima between the sexual functions, or alternatively, an outcrossing advantage (Charlesworth 1999).

Sex chromosomes are one of the potential determinants of sex. Pairs of sex chromosomes control sex at the individual level, and differ from autosomes mainly in their partial loss of meiotic recombination, and because one of them is limited to one of the sexes (Charlesworth 2016). Some sex chromosome pairs are heteromorphic in karyotypes, while others are homomorphic. Few plant sex-determination systems and sex chromosomes have been studied in detail (Ming et al. 2011), and even fewer of these originated independently. This severely limits comparative studies aiming to understand the incidence and stability of sex chromosomes in the tree of life (The Tree of Sex Consortium 2014) and the identification of universal patterns in their evolution and structure. Beyond fundamental evolutionary questions, knowledge of sex-determination systems also has important applications for example in molecular gender phenotyping of juvenile plants in agriculture, plant breeding, and conservation.

This study aimed to develop and apply a novel and robust method to characterise sex-determination systems, unravel the sex-determination system of *Nepenthes* pitcher plants, and investigate biological questions related to the origin of dioecy in this genus.

### Study system

*Nepenthes* (Nepenthaceae, Caryophyllales) comprises c. 140 taxa of perennial vines and shrubs (Cheek & Jebb 2001; McPherson 2009) occurring mostly in Southeast Asia (Clarke 1997). All species are carnivorous plants which supplement their nutrient budget by killing and digesting insects (among other prey), enhancing growth and flowering (Pavlovič & Saganová 2015; Moran & Moran 1998). The complex physiology of carnivory takes place in modified, jug-shaped leaves called pitchers (Juniper et al. 1989; Moran & Clarke 2010).

All *Nepenthes* are dioecious and hermaphrodites are not documented, while the closest relatives, families Ancistrocladaceae, Dioncophyllaceae, Droseraceae, and Drosophyllaceae (Cuénoud et al. 2002; Renner & Specht 2011) are entirely hermaphroditic. Sexual dimorphism in *Nepenthes* could be restricted to the reproductive structures (Kaul 1982), but other traits lack study. Male and female flowers (Figure 1a) are highly diverged because alternative reproductive organs abort early in development (Subramanyam & Narayana 1971). Inflorescences generally share the same structure in males and females (raceme, panicle), but may differ in tepal colouration and shape, peduncle and rachis length (reviewed in Supporting Information (SI) Table S1; Clarke, 1997, 2001, McPherson, 2009, 2011; Clarke *et al*., 2011; described for only 46 of 138 taxa), and nectar production (Frazier 2001; Kato 1993). Male inflorescences bear in general more flowers (Frazier 2001). Sexual dimorphism in ecology (Barrett & Hough 2013) may exist in *Nepenthes*: In disturbed habitats, adult *N. gracilis* and *N. rafflesiana* were strongly male-biased under open canopy (83% and 100% male individuals, respectively) but were slightly female-biased under closed canopy (34% and 40% males; Frazier 2001), consistent with the hypothesis that males tolerate drier and hotter conditions.

**Fig. 1.**
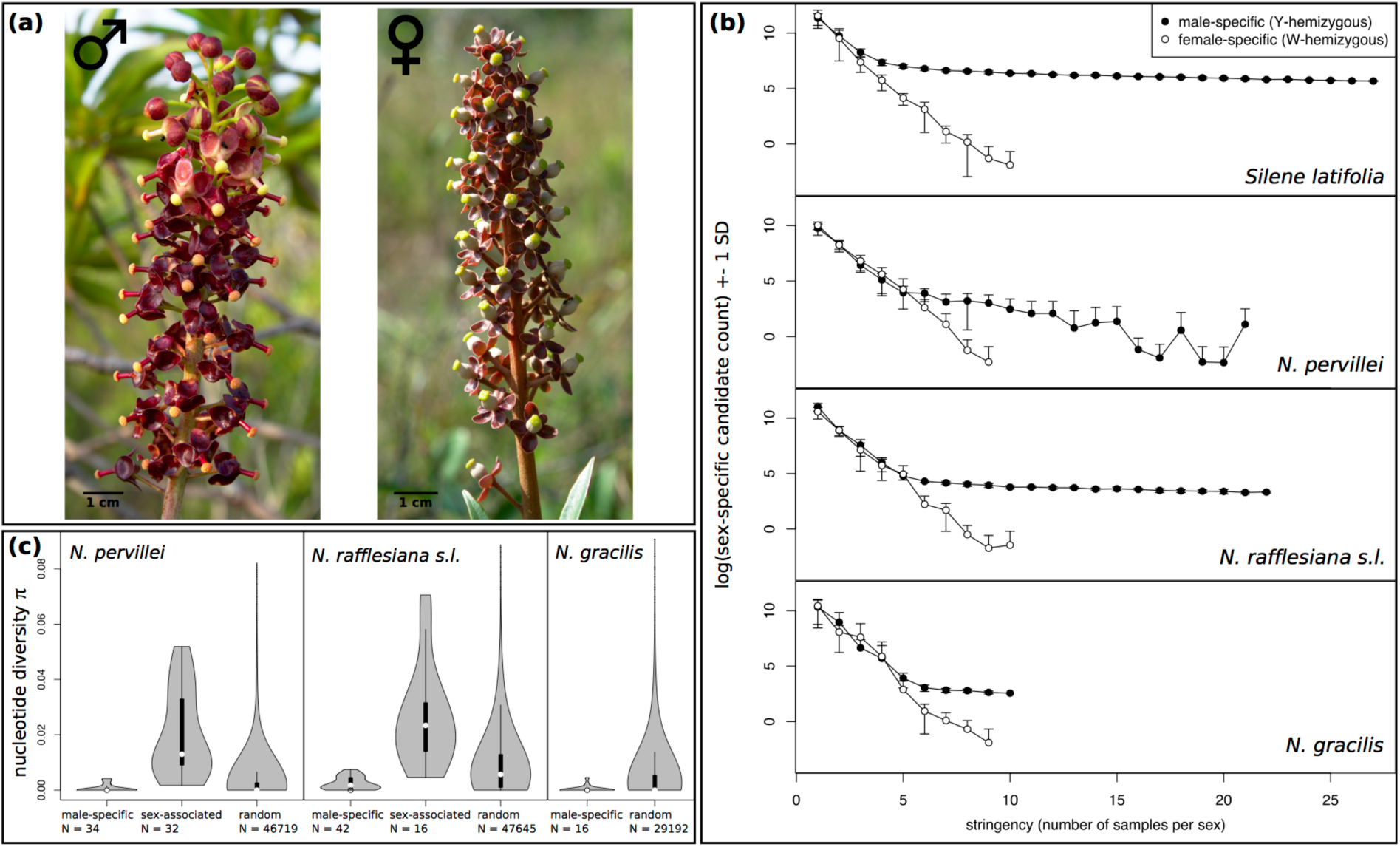
(a) male inflorescence of *N. rafflesiana s.l*. (left) and female inflorescence of *N. mirabilis* var. *globosa* (right). (b) Evidence for male-specific markers and an XY sex-determination system in *Silene latifolia* and three *Nepenthes* spp. (privacy rarefaction curves). Shown are counts of sex-specific markers (y-axis) as a function of the number of samples per sex used to score sex-specificity (x-axis). Sex-specific markers are defined as RAD-tags with mapped reads in all samples from one sex but without any mapped reads in the same number of samples from the other sex. Dots represent averages and whiskers one standard deviation of 200 bootstrapped combinations of males and females. Note natural log-scale of y-axis and hence undefined negative values in SD ranges. (c) Mean per-site nucleotide diversity n of RAD-tags in male *Nepenthes* of three taxa for male-specific, sex-associated, and random RAD-tags. All RAD-tags mapping 3–75 reads in >=75% of males per population were included. The same sets of individuals are considered in each category. Sex-associated RAD-tags were absent in *N. gracilis*. Median = white dot, box = 25%−75% quartiles, whiskers = 1.5*interquartile range, violin = estimated kernel density.

The sex-determination mechanism in *Nepenthes* is unknown, but there are no reports of plasticity or reversal of the gender in nature or cultivation (Clarke 2001), suggesting stable sex determination during early development, or a genetic basis. Heteromorphic sex chromosomes are unlikely since a wide range of species share indistinguishably small and uniform chromosomes (2n=80; Heubl & Wistuba 1997).

### Analysis of sex-determination systems

Cytogenetics and linkage analysis in families are traditional methods to study sex-determination and sex linkage of genes (Charlesworth & Mank 2010). However, these strategies fail in many dioecious organisms because of uninformative karyotypes and prohibitive logistics of breeding. Here we avoid cytology and controlled crosses by instead scanning natural populations for associations between sex and genetic markers.

Two main categories describe molecular genetic sex differences (sex-linkage): sex-association and sex-specificity. Sex-associated loci differ in allele frequency between sexes, and either directly determine sex, for example in polygenic sex-determination systems (reviewed in Bachtrog et al. 2014), or display linkage disequilibrium (LD) with sex-determining loci. Sex-specific loci are private to one sex and entirely absent from the other. They indicate partial divergence between male and female genomes, likely following recombination suppression around sex-determining loci (Charlesworth 1991). Pairs of sex chromosomes can be classified as either heteromorphic or homomorphic. In the former, cytogenetic (optical) techniques indicate chromosome divergence in size or structure, whereas in the latter the differences are so subtle that molecular genetic methods are required to resolve them. Male-limitation is referred to as XY-heterogamety and female-limitation as ZW-heterogamety. Sex-specific regions of Y or W chromosomes occur as single copies (haploid, hemizygous) in diploid tissue.

Genomes of both natural and cultivated populations may be scanned for sex differences. Candidate sex-associated or sex-specific loci are expected to display strong mutual LD, and reduced genetic diversity relative to autosomal regions (Wilson Sayres et al. 2014). Lack of candidate loci may indicate either non-genetic (environmental) sex-determination, a very small sex-specific region, or multi-locus sex-determination. To date, few studies have used population genetics to unravel sex-determination systems and identify sex-linked loci without pedigrees. Indeed, chances to find a potentially small sex-linked region with few markers in a large genome are poor. Noteworthy exceptions include the discovery of a sex-specific microsatellite locus in European tree frogs, revealing males as heterogametic in this species (Berset-Brändli et al. 2006). However, marker availability is no longer limiting since the development of restriction site associated DNA sequencing (“RAD-seq” and related methods; Elshire et al. 2011; Peterson et al. 2012; Baird et al. 2008). These methods successfully identified sex-linked markers and sex-determination systems without pedigrees in diverse organisms such as Crustaceans (Carmichael et al. 2013), *Anolis* lizards (Gamble & Zarkower 2014), and moore frogs (Brelsford et al. 2017).

Several studies have reported apparent sex-specific loci in both sexes (Gamble & Zarkower 2014; Bewick et al. 2013; Heikrujam et al. 2015; Brelsford et al. 2017). However, it is generally expected that there exists only one sex-specific chromosome and only one heterogametic sex. Even in the unusual XYW system (Orzack et al. 1980) or XY-ZW transitional phases (Sander van Doorn & Kirkpatrick 2010), only one sex has a private (dominant) sex chromosome. The paradoxical reports may represent false positives, a problem exacerbated in RAD-seq data with their highly stochastic genotype presence–absence polymorphisms (Mastretta-Yanes et al. 2015). As noted by previous authors (Gamble & Zarkower 2014; Gamble et al. 2015), the rate of false positive sex-specific loci decreases the more males and females are compared, but true positives may be lost at an even higher rate. There exists so far no solution to this quality–quantity trade-off.

### Aims of the study

We investigated the unknown sex-determination system of dioecious *Nepenthes* pitcher plants and examined whether sex chromosomes across their radiation are derived from a single pair of ancestral autosomes. We developed a statistical procedure that allows distinguishing between true sex-specificity and stochastic absence, and applied this approach to *Nepenthes*. Specifically, we addressed the following questions: (1) Are there sex-specific and sex-associated RAD-tags in *Nepenthes*? (2) Do sex-specific and sex-associated RAD-tags conform to theoretical population genetic expectations? (3) Are sex-specific and sex-associated regions shared among different *Nepenthes* species? (4) Do these regions contain expressed genes that may contribute to differences between the sexes? Based on our results we further developed a molecular sexing assay for *Nepenthes*. This tool is suitable for identifying the sex of juveniles and non-flowering adult plants in ecological research, conservation and horticulture.

## Materials and Methods

### Sampling, ddRAD-seq and genotyping

Samples genotyped by ddRAD-seq (Peterson et al. 2012) and used for this study are shared with those used by Scharmann *et al* (in revision), and we refer readers to this publication for specific details. In brief, natural populations of *Nepenthes* were sampled in Brunei Darussalam (Borneo), Singapore, and the Seychelles. Genomes were Illumina-sequenced in strongly reduced form by focussing on DNA restriction fragments (two enzymes). RAD-tags (contigs) were *de novo* assembled by clustering reads, followed by mapping, genotype calling and quality filtering. Scans for sex-specific and sex-associated markers were conducted separately on three taxa: *N. pervillei* Blume (28 males and 22 females from six populations on Mahé), *N. gracilis* Korth. (ten males and ten females from one population in Brunei), and *N. rafflesiana sensu lato* (in total 39 males and 22 females combined from four populations). We here define *N. rafflesiana s.l*. as *N. rafflesiana* Jack from Singapore (ten males, four females) and the Bornean entities *N. rafflesiana* “typical form” (twelve males, six females), *N. rafflesiana* “giant form” (Clarke 1992, 1997; five males, three females) and *N. hemsleyana* Macfarlane (Scharmann & Grafe 2013; eleven males, eight females).

The sex of individual was established on the basis of fresh or dry inflorescences, or else molecularly using a preliminary sexing assay for *N. rafflesiana s.l*. (SI Methods S1). To increase the phylogenetic range of our study and to validate molecular sexing, we furthermore included individuals of known sex for additional species from the field, and species that flowered in cultivation (table in SI Methods S4). Fresh leaf material was stored in a nucleic acid preserving buffer (Camacho-Sanchez et al. 2013) until further use.

To validate our method for detection of sex-specific markers, we also genotyped several individuals of *Silene latifolia* Poiret, a species with a known XY sex-determination system and heteromorphic sex chromosomes. Details of *Silene* sampling and genotyping are provided in SI Methods S2.

### Detection of sex-specific RAD-tags

We define sex-specific RAD-tags as mapping sequencing reads from one of the sexes exclusively. The number and identity of sex-specific RAD-tags both carry uncertainties because they depend on the number and identity of male and female individuals compared. We evaluated these uncertainties by resampling methods, separately for each of three *Nepenthes* taxa and *Silene latifolia*. Sex-specificity was tested quantitatively, i.e. for deviation of the observed number of sex-specific RAD-tags from zero, and qualitatively for each RAD-tag.

Unbiased comparisons among data sets with different numbers of individuals and sex ratios were achieved by subsampling male and female individuals such that a 1:1 sex ratio was maintained. To capture the uncertainty over different combinations of individuals, we bootstrapped without replacement for each possible subsample size 200 random sets of males and females. In the quantitative tests, we counted sex-specific RAD-tags for each combination. This generated two distributions, the bootstrapped observed male- and female-specific RAD-tag counts. Then, these two distributions were separately compared to a null distribution derived from permutations of the sexes. The null distribution estimates how many RAD-tags would appear to be sex-specific if the individuals were interchangeable, i.e. if there were no true sex-specific RAD-tags. We calculated a *p*-value as the proportion of permuted sex-specific counts equal to or larger than the mean of the observed male- resp. female-specific count distribution.

The qualitative test assessed the confidence in sex-specificity for each locus. We again bootstrapped 200 random sets of males and females without replacement from the available individuals, for each possible subsample size. Then, we counted for each locus in how many of the bootstrapped male–female comparisons it emerged as sex-specific. True sex-specific RAD-tags are expected to appear more frequently in such comparisons than false positives whose occurrence is random. The bootstrap support value for sex-specificity is the count divided by the number of bootstraps. We only reported RAD-tags with 50% or higher bootstrap support.

We named this algorithm privacy rarefaction and implemented it in a multithreading python script that calls samtools (Li et al. 2009) to read mapping data from .bam alignments (will be available at https://github.com/mscharmann/).

### Detection of sex-associated SNPs

We conducted chi-squared tests on frequency counts for all bi-allelic SNPs versus sex in PLINK v.1.07 (Purcell et al. 2007). Maximum 25% absent genotypes were tolerated per SNP, and candidate SNPs were accepted as sex-associated at a false discovery rate (Benjamini & Hochberg 1995) smaller or equal to 0.05.

### Population genetics of candidate RAD-tags

We tested whether LD in male populations differed between sex-specific resp. sex-associated SNPs and the genomic average, represented by 100 randomly selected SNPs. LD (r^2^) was calculated between but not within RAD-tags using VCFtools v0.1.15 (Danecek et al. 2011). The same contrasts were made for nucleotide diversity π, which was averaged per RAD-tag using SNPs from .vcf genotypes (VCFtools) while the total number of observed sites per RAD-tag was taken from .bam alignments, applying the same filters to both data (minimum read depth 3, maximum read depth 75, maximum genotype absence 0.25). Significance of differences was evaluated by a randomisation test in R (bootstrap resampling from observed data at equal sample sizes, permutation of observations; *p*-value = proportion of resampled datasets with difference in means greater or equal to the observed difference; 100k replicates).

### Comparison of candidate RAD-tags to a male inflorescence transcriptome

We sequenced and assembled the transcriptome of a developing male inflorescence of *Nepenthes khasiana* Hook.f. (SI Methods S3) to identify and annotate sex-linked candidate loci. Fresh inflorescences of the species used for ddRAD-seq were not available in cultivation. The transcriptome was searched (a) by BLAST for similarity to candidate RAD-tags (thresholds >= 90 aligned bases and >= 75% identity), and (b) by repeating privacy rarefaction with ddRAD-seq reads directly mapped to the transcriptome rather than the ddRAD-seq assembly (bwa mem; Li 2013; not filtering mapping quality, allowing multiple mappings).

Candidate transcripts from both approaches were annotated by BLAST search against NCBI Genbank nt (version as of Nov 7, 2016), and NCBI Genbank nr (version as of Mar 26, 2016). Transposable elements were detected using RepeatMasker 4.0.6 (Smit et al. 2013) v. 20150807 (Eukaryota). Proteins with at least 50 amino acids were predicted by TransDecoder (Trinity package) and annotated against NCBI Genbank nr, UniProt Swiss-Prot (version as of Aug 17, 2016), and *Arabidopsis thaliana* proteins in UniProtKB (version as of Apr 3, 2016). PFAM domains were detected using hmmer 3.1b1 (Eddy et al.). Database hits were accepted at e-value <= 10^−5^.

### PCR validation

Candidate sex-specific RAD-tags were chosen for PCR validation based on a ranking of the highest stringency level reached, bootstrap support, taxonomic overlap, and the quality of annotation of matching transcripts. PCR primers were designed in Geneious R6 (Biomatters Ltd., Auckland, New Zealand). PCR reactions were performed in 15 *μ*l volumes containing 2.5 mM MgCl_2_, 250 *μ*M of each dNTP, 0.375 units of GoTaq DNA polymerase (Promega, Wisconsin, USA), 1x GoTaq Flexi buffer (Promega), 0.5 *μ*M of each primer, and 1 *μ*l of template DNA (2–20 ng/*μ*l). After initial denaturation for 2 min at 95°C, 30 cycles were run with denaturation at 95°C for 30s, annealing at 50°C for 30s, and extension at 72°C for 1 min, followed by a final extension step of 5 min at 72°C (Labcycler, SensoQuest, Göttingen, Germany). PCR products (2 *μ*l) were separated by electrophoresis in a 2% agarose gel and visualised through fluorescent staining (GelRed, Biotium Inc., Hayward, CA, USA).

## Results

### Sex-specific RAD-tags

Qualitatively consistent signatures of male-specific RAD-tags were detected independently in *N. pervillei, N. gracilis, N. rafflesiana s.l*., and *Silene latifolia* (Figure 1b). Numbers of candidate RAD-tags decreased monotonically with subsampling size. This drop in number of shared RAD-tags with increasing number of samples is an inherent and typical property of RAD-seq data, which have large genotype absence caused by a combination of restriction site mutations and stochasticity during library preparation, Illumina sequencing, and bioinformatics (reviewed in Mastretta-Yanes et al. 2015).

The evolutionary plant model for dioecy associated with an XY sex-determination system, *Silene latifolia*, constitutes a positive control for the identification of male-specific RAD-tags since it contains a well-differentiated Y-chromosome. Simultaneously, *S. latifolia* provides a negative control for a ZW-system as evidenced by the decay of false positive female-specific RAD-tags with increasing stringency (subsample size) in all four taxa.

### Sex-associated SNPs

We detected bi-allelic SNPs associated with the phenotypic sex in *N. pervillei* and *N. rafflesiana s.l*., as well as in *Silene latifolia*, but not in *N. gracilis*. Almost all sex-associated SNPs had an allele frequency close to 0.5 and near-complete heterozygosity in males, but were close to fixation and thus homozygous in females (SI Table S2). The proportion of sex-associated bi-allelic SNPs identified was much lower in *Nepenthes* (*N. pervillei*: 97/38,783=0.25%; *N. rafflesiana* s.l.: 37/222,188=0.017%; *N. gracilis:* 0/50,483=0%) than in *S. latifolia* (2,376/149,311=1.6%).

### LD among sex-specific RAD-tags and sex-associated SNPs

Sex-specific and sex-associated genomic regions are expected to experience little or no recombination, which should lead to increased LD. Contrary to expectations, LD among sex-specific RAD-tags did not differ from the genomic background (*p* = 0.74) in *N. pervillei*, whereas mean LD among SNPs in sex-associated RAD-tags was elevated by 0.077 units over the genomic background (*p* <= 10^−5^). In *N. rafflesiana s.l*., mean pairwise r^2^ among SNPs located in sex-specific RAD-tags was 0.1 units higher than in the genomic background (*p* <= 10^−5^), whereas LD among sex-associated RAD-tags was only slightly higher than the genomic background (c. 0.004 units, *p* = 0.0011). These tests could not be conducted for *N. gracilis* because no sex-associated RAD-tags were identified and only two sex-specific RAD-tags contained SNPs (r^2^ = 0.15625).

### Nucleotide diversity of sex-linked RAD-tags

Mean π in male-specific RAD-tags tended to be lower than the genomic background in all three taxa (Figure 1c). This difference was significant for *N. rafflesiana s.l*. (*p* = 0.0005), but not for *N. pervillei* and *N. gracilis* (*p* = 0.11 and *p* = 0.065, respectively). In contrast, mean *π* in sex-associated RAD-tags in males was increased over the genomic background for both *N. pervillei* (*p* <= 10^−5^, Figure 1c) and *N. rafflesiana s.l*. (*p* = 0.0043; Figure 1c).

### Shared candidate loci between species, functional annotations and PCR validation

We recovered six shared candidate sex-specific RAD-tags at stringency level >= 5 in both *N. gracilis* and *N. rafflesiana s.l*., whereas *N. pervillei* shared no candidates with either (SI Table S3). There was no overlap in sex-associated SNPs between the *Nepenthes* species, and no direct overlap between sex-specific RAD-tags and those with sex-associated SNPs. However, one male-specific RAD-tag of *N. gracilis* and one RAD-tag with sex-associated SNPs of *N. rafflesiana s.l*. both matched with high confidence to the same inflorescence transcript containing a DUF4283 (domain of unknown function, http://pfam.xfam.org/family/PF14111, 09.11.2016).

One male-specific RAD-tag of *N. pervillei* aligned to the transcript of a bHLH transcription factor, and the best matches in all accessed databases were consistently to predicted orthologs of the *Arabidopsis* gene DYSFUNCTIONAL TAPETUM 1 (DYT1). A further sex-associated RAD-tag of *N. pervillei* matched a transcript annotating as *A. thaliana* SEPALLATA-1 (SEP1). This RAD-tag aligned to the predicted 3’-UTR of the putative SEP1-ortholog, and contained two SNPs which were both homozygous in 95% of females and heterozygous in 96% of males. However, further comparisons of putative X–Y divergence of SEP1 were not possible because the male inflorescence transcriptome reads were not heterozygous. In *N. gracilis*, a male-specific RAD-tag matched a long transcript similar to a mitochondrial NADH-ubiquinone oxidoreductase from *Beta vulgaris* (Swiss-Prot). All further candidate loci contained either traces of transposable elements, or no known sequence motifs (SI Table S3).

Complementary to the sex-specificity scan on the ddRAD-de *novo* reference, we repeated privacy rarefaction by directly mapping the ddRAD reads to the male inflorescence transcriptome, with the aim to recover further annotated candidate genes. This approach identified seven transcripts as male-specific in at least one species (SI Table S3). We considered only high-confidence male-specific candidate transcripts, reaching at least stringency level four and bootstrap support greater 0.5 in at least one species. No female-specific transcripts (false positives) reached this stringency. A single transcript was male-specific in *N. rafflesiana s.l*. but could not be annotated. Four close transcript “isoforms” (Trinity assembler) were male-specific in both *N. gracilis* and *N. rafflesiana s.l*., but they lacked similarity to any known motif except for one isoform similar to a Jockey-1_Drh retrotransposon. However, two transcripts were male-specific in both *N. pervillei* and *N. rafflesiana s.l*., and one of these also matched a *N. pervillei* male-specific RAD-tag (see above). These two transcripts appear to be close isoforms (putative intron presence–absence), and both annotated as DYT1 (see above).

We tested by PCR whether the putative DYT1-ortholog is male-specific in a broad range of *Nepenthes* species. A single PCR product of approximately 290 bp length was observed exclusively and consistently in phenotypically sexed male *Nepenthes* but never in females (SI Methods S4). We tested multiple males and females (1 vs. 3 to 3 vs. 3) for eight taxa, and 1–2 individuals from 14 further taxa. Presence–absence of the PCR product was fully consistent with the phenotypic sex of all tested individuals (n=56). Sanger sequencing of the PCR product confirmed the identity of the target region. Hence, this locus is male-specific across a phylogenetically broad range of *Nepenthes* species and can be used for molecular sexing of individuals.

## Discussion

### The *Nepenthes* sex-determination system

In natural populations of *Nepenthes* we discovered both sex-associated markers that were predominantly heterozygous and displayed high nucleotide diversity in males but were mostly homozygous in females, as well as multiple male-specific markers displaying elevated LD and reduced nucleotide diversity. The latter is consistent with theoretical expectations for sex-specific loci, which are hemiploid and have only 1/4 (assuming equal sex ratio) of the effective population size compared to the autosomal genome. Both patterns can arise and persist in interbreeding populations as a consequence of physical linkage to sex-determining loci, (partial) cessation of recombination in the MSY, and X–Y divergence. Together, these findings reveal a genetic basis of sex-determination in *Nepenthes* spp. in which males are the heterogametic sex, i.e. an XY-system. Our interpretations are supported by congruent patterns inferred in the well-known XY-heterogametic *Silene latifolia*.

### Method to extract sex-specific loci from population genotype data

Methods to rapidly genotype individuals across the genome, such as RAPDs, AFLPs, and more recently RNA-seq and ddRAD-seq were repeatedly used to identify sex-specific markers. All these approaches suffer from the common problem that marker absence in some individuals (e.g. due to polymorphism, low coverage or technical artefacts) must be distinguished from true marker absence in the entire sex. Erroneous inference of marker absence in one sex leads to false positive sex-specific markers for the other sex. This likely played a role in studies reporting both male- and female-specific markers in single populations (Gamble & Zarkower 2014; Bewick et al. 2013; Heikrujam et al. 2015; Brelsford et al. 2017).

Theoretical expectations for fully dioecious diploids in quasi-panmixia imply that a population can not harbour both male- and female specific loci at the same time. Under these assumptions, at most one sex, or else none, carries sex-specific alleles or loci as derived from the principles of diploid inheritance (Mendel 1866): If sex is determined by a single locus, here not understood as a single gene but more broadly as a non-recombining DNA sequence that may contain any number of genes, it must follow dominant-recessive inheritance (where absence constitutes a recessive allele). Co-dominance can be excluded because it would produce hermaphrodites or steriles, violating the assumption of a fully dioecious population. Consequently, all loci that are not perfectly physically linked to the sex-determining locus are expected to be shared by both sexes, and sex-specific loci and alleles must be located in a single, non-recombining genomic block that includes a dominant sex-determining allele. If sex is controlled by more than one locus, as in quantitative or polygenic sex-determination, however, by definition no single locus or allele controls sex, and hence all loci and alleles are expected to be shared by both sexes.

We argue that erroneous inference of marker absence, and thus false positive identification of sex-specific markers, results from insufficient consideration of uncertainty in presence-absence within and between sexes. This includes sex bias in both sample size and genetic structure of the screened population. We eliminated this problem through replicated downsampling from a larger pool of observations (individuals) in the same way as rarefaction in community ecology eliminates sampling bias when comparing species richness (Gotelli & Colwell 2001). However, instead of the resampled counts in two groups (habitats), we record the identity and level of sharing (or privacy) between groups (sexes), which is not of interest in conventional rarefaction analysis. As a result, privacy rarefaction curves, as we name them here, decay rather than increase towards a plateau with increasing sub-sample size (=stringency). An empirical statistical solution to differentiate between random and true privacy of genomic loci, like the one detailed here, has to our knowledge not been applied before (but see Schlüter & Harris 2006; Szpiech et al. 2008; Kalinowski 2004 for applications in genetic fingerprinting and diversity estimation).

Previous approaches to detect sex-specific loci from wild populations sequenced several males and females and then scored which loci are absent from all males and present in all females, and vice-versa. Naïvely, the scoring is done once with all individuals together, frequently with different sample sizes for males and females. Although biologically plausible results are sometimes obtained (e.g Utsunomia et al. 2017), many studies report sex-specific markers for both sexes (Gamble & Zarkower 2014; Bewick et al. 2013; Heikrujam et al. 2015; Brelsford et al. 2017). These results evidently contain artefacts because they do not conform to the Mendelian expectation that only one of the sexes can harbour sex-specific loci. Thus, results for one sex are exclusively false positives, whereas results for the other sex may encompass both true and false positives. To distinguish true from false positives, researchers then typically attempt to validate many candidate loci by PCR in reference populations. Here, we report that the problem can partially be solved in-silico by our novel privacy rarefaction algorithm. It removes false-positives and assigns a quality score to each candidate locus, thus greatly reducing the number of candidate sex-specific markers to validate.

Our use of privacy rarefaction for the inference of sex-specificity aims at identifying the most likely true-positive candidates in noisy datasets, at the possible cost of a high false-negative rate. Privacy rarefaction curves are illustrated in Fig. 1. In these, false-positive candidates are evident: If apparent sex-specific loci are identified in both sexes (typically at low stringency), then false-positives are present in at least one of the sexes. This happens when too few males and females are compared, e.g. at two males versus two females we typically obtained tens of thousands of sex-specific candidates, most of which were false-positives. These false-positives appeared sex-specific because they were not present in the dataset in all individuals, due to technical artifacts, low coverage or molecular polymorphisms, and they by chance coincided with sex. Thus, different combinations of few males and few females yield large but inconsistent sets of sex-specific candidates, most of which will be false-positives. With increasing numbers of males and females, false-positives are progressively eliminated as the collective male-resp. female sets of loci approach the real genomic composition of males resp. females. Different combinations of many males and many females yield small but largely consistent sets of sex-specific candidates. This happened in our experience from c. ten males and females upwards, and our script reports this concistency as a bootstrap support value. The dropout of false-positives characterises the initial steep decay of the privacy rarefaction curves and continues until only one sex contains specific loci. This is a critical point which is diagnostic for the heterogametic sex. We expect that the sex-specific candidates that occur at and beyond this stringency have a false-positive rate near zero, because there are no more false-positives for the homogametic sex. However, at this point we expect to have a high false-negative rate that arises from stochastic absence. True sex-specific loci are classified as not sex-specific if they are absent in a subset of the investigated individuals of the respective sex. The slower decay of privacy rarefaction curves at higher stringencies is due to the dropout of true-positive candidates.

As the noisy individual presence-absence is smoothed by aggregation over many individuals, one may argue that sex-specificity should be scored only for the maximum possible number of males and females in a dataset. However, this strategy can miss true sex-specific loci entirely if the data are noisy and true sex-specific loci have inconsistent and low coverage, e.g. when large genomes with very small sex-specific regions are sequenced at low depth. But if combinations of fewer males and females are also evaluated, it is possible to find that one sex consistently yields more candidates than the other, which indicates that true positives exist for that sex. Our script calculates a p-value for the difference between male- and female-specific candidate counts. Importantly, the privacy rarefaction algorithm per se can not affect the false-negative rate because it is not based on statistical model assumptions - false-negatives are given by stochasticity in read presence-absence, i.e. due to the sampling design, wetlab and in-silico procedures.

A recent review discussed six available methods for identification of sex-linked sequences (Muyle et al. 2017). Our method, the combination of population genomic data (individual data generated by any sequencing method, e.g. RNA-seq, reduced-representation libraries, whole-genome) with privacy rarefaction, fills a gap in these existing methods, most importantly because it does not depend on breeding or an assembled reference genome. Some of these methods further require prior knowledge of the heterogametic sex and exhaustive genome sequencing of males and females. Our method is related to the Bayesian classification algorithm detsex (Gautier 2014) in some of its scope and the required input data. However, privacy rarefaction is model-free, was here used successful with less than half of the individuals recommended for detsex (i.e. 10–20 per sex as opposed to >40; Muyle et al. 2017), and copes with very large and noisy datasets with high missingness. Privacy rarefaction curves thus offer a unique, simple and robust way to judge whether true sex-specific loci exist and which sex is heterogametic. The established methods for identification of sex-linked sequences are more suitable for projects that aim to further study previously identified sex chromosomes, while our approach is efficient in the first phase of investigation of new species, i.e. rapid de novo-discovery of unknown sex determination systems and cytogenetically homomorphic sex chromosomes with some of their basic properties. But privacy rarefaction can also complement more advanced projects, because it identifies sex-specific loci (Y-resp. W-specific loci), a major class of sex-linkage that is neglected by family segregation analyses due to the lack of recombination.

Based on available experience, we expect the power of our method to discover sex-specific markers and heterogamety to be limited by the true number of sex-specific loci, the design of population sampling, and the relationship of stochastic noise in genotype presence (consistency of library preparation and read coverage) to the number of individuals per sex. The method performs better the more truly sex-specific loci exist, the higher the read coverage, the more individuals of each sex are included, and the less the population deviates from panmixia (both family structure and geographic structure should be avoided). For example, in a species with XY sex-determination, full-sibling males and females do not contain identical X-chromosomes if their parents carried polymorphic X chromosomes; this may lead to the apparent paradox of finding both male- and female-specific markers. However, this was not the case in our analysis of three *Nepenthes* species and *S. latifolia*, in which female-specific loci were significantly outnumbered by male-specific loci at stringency levels greater than six, and fell to zero above stringency levels ten or eleven (Figure 1b). Moreover, our method is robust to modest levels of erroneous gender phenotyping. We consequently expect privacy rarefaction to be useful for diverse study systems.

### Properties of the *Nepenthes* XY system

*Nepenthes* karyotypes suggest that the sex chromosomes are homomorphic (Heubl & Wistuba 1997), matching the relatively lower proportion of Y-specific and sex-associated RAD-tags in *Nepenthes* compared to *S. latifolia* with its disproportionally large Y-chromosome. Based on theoretical expectations (above), we interpret the occurrence of true sex-specific markers as evidence for a single sex-determining genomic region.

An expressed gene homologous to *Arabidopsis* DYT1 was found to be male-specific in both *N. pervillei* and *N. rafflesiana s.l*. PCR tests confirmed that this gene is exclusively amplified in males, but never in females, of 22 *Nepenthes* species. We interpret this result as evidence for a conserved core of the *Nepenthes* MSY, given that *N. pervillei* and *N. rafflesiana s.l*. span the largest phylogenetic distance in the genus (Meimberg et al. 2001; Mullins 2000). This shared core MSY further suggests a single common origin of dioecy in *Nepenthes*. Dioecy most likely evolved after the split of dioecious Nepenthaceae and hermaphroditic Droseraceae from their most recent common ancestor (MRCA) at 71.1 (CI 44.2-98.0) Mya, and before the MRCA of extant *Nepenthes* at 17.7 (CI 11.0-24.3) Mya (SI Methods S5). *Nepenthes* does not conform to the view that sex chromosome pairs increasingly diverge over time (Bachtrog et al. 2014), but rather fall into the category of lineages with old and homomorphic sex chromosomes (e.g. ratite birds: Vicoso et al. 2013, brown algae: Ahmed et al. 2014).

During the radiation of *Nepenthes*, the MSY has diverged between species. Only six out of 135 male-specific RAD-tags were shared between *N. rafflesiana s.l*. and *N. gracilis*, and none were shared with the more distant *N. pervillei*. Male-specific loci shared between *N. pervillei* and *N. rafflesiana s.l*. were only recovered by directly mapping ddRAD reads to the male inflorescence transcriptome. This suggests that absence of shared sex-specific RAD-tags should not be interpreted as evidence for independent origins of sex chromosomes. Further evidence for a common origin but subsequent interspecific divergence of sex chromosomes is found in a DUF4283-transcript, which was male-specific in *N. gracilis* but sex-associated in *N. rafflesiana s.l*. Apparently, X and MSY alleles (i.e. gametologs) have lost homology (threshold 90% identity) in the former but not in the latter species.

Apart from sequence divergence, variation in absolute and relative abundance of male-specific and sex-associated markers is consistent with variation in the size of the MSY among species, as may be expected in independent evolutionary lineages. Alternatively, limited overlap in male-specific RAD-tags among species outside a core sex-determining region may be compatible with sex chromosome turnover, where the sex-determining region translocated into different chromosomal backgrounds (Blaser et al. 2014). Until the genomes of multiple species have been sequenced and compared, this alternative cannot be fully excluded. Yet this aspect does not affect our conclusion of a single common origin of the MSY and dioecy in *Nepenthes*.

### Non-coding DNA and special significance of DYT1 and SEP1

Of the 38 sex-linked inflorescence transcripts (identified by 41 matching sex-linked RAD-tags), 34 (89%) could not be annotated or contained transposable elements (TEs). Thus, the great majority of sex-linked genomic regions retrieved from *Nepenthes* correspond to non-coding sequences and TEs, and here they were more common than in non-sex-linked transcripts (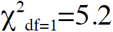, p=0.02). Nine out of 13 sex-linked TE-transcripts belonged to a particular family of TEs, the *gypsy*-like retrotransposons, whereas this family was much less common among non-sex-linked TE transcripts (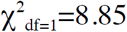, p=0.003). TE accumulation is expected in non-recombining regions, and has been reported in species with both heteromorphic (e.g. *Silene*, Čermák et al. 2008) and homomorphic sex chromosomes (e.g. Papaya, Wang et al. 2012). The match of a sex-associated RAD-tag in *N. gracilis* to a mitochondrial NADH-ubiquinone oxidoreductase was unexpected but may represent either an unspecific match of the short (96 bp) RAD-tag to the inflorescence transcript, or else a cyto-nuclear transfer to the sex chromosomes. The occurrence of organellar genes on plant sex chromosomes has been documented in other species (Steflova et al. 2013).

Two expressed sex-linked genes are of special interest. First, we identified a *Nepenthes* homolog of DYT1 as a genus-wide male-specific (MSY) locus. DYT1 is essential for tapetum development and thus pollen fertility in *A. thaliana* (Zhang et al. 2006), rice (Cai et al. 2015; Jung et al. 2005; Wilson & Zhang 2009), and tomato (Jeong et al. 2014), and we speculate that *Nepenthes* DYT1 is also functionally conserved. Direct validation in *Nepenthes* is currently not feasible due to the lack of transformation protocols and the long generation times. Our analysis suggests that *Nepenthes* DYT1 is absent from females and thus absent from the X chromosome. Such a deletion of DYT1 from the X chromosome would constitute a recessive male-sterility mutation, which is required early in the evolution of dioecy for the transition from a hermaphroditic to a gynodioecious mating system (Charlesworth & Charlesworth 1978).

The second gene of interest is a *Nepenthes* homolog of the homeotic MADS box gene SEP1, an early-acting regulator of floral organ identity in *Arabidopsis* (Pelaz et al. 2000), which was sex-associated in *N. pervillei*. There was near-perfect heterozygosity in two SEP1-linked SNPs in males, whereas these positions were highly homozygous in females, consistent with a location in little- or non-recombining regions of the X and Y chromosomes and gametolog sequence divergence. If SEP1 is functionally conserved in *Nepenthes*, a major component of floral organ development pathways (Theißen et al. 2016) is located on *Nepenthes* sex chromosomes. We thus hypothesize that SEP1 is involved in unisexual flower development in *Nepenthes*. In *Silene latifolia*, however, SEP1 homologs are not directly involved in sex-determination and not located on the sex chromosomes (Matsunaga et al. 2004).

The possible roles of DYT1 and SEP1 in the origin of dioecy in *Nepenthes* require further attention. Even if these genes do not directly determine sex in extant *Nepenthes*, they may have been involved in sexually antagonistic selection during the evolution of dioecy because a deletion of DYT1 or alternative SEP1 alleles might abort non-functional organs at an earlier developmental stage, thus saving resources. However, the completely unisexual floral morphology of extant *Nepenthes* implies that further genetic differences exist between males and females.

### Ecological causes of dioecy in *Nepenthes*

In general, selection for an outcrossing mating system and sexual conflict are hypothesized to drive transitions from hermaphroditism to dioecy (Charlesworth 1999). The *Drosera* species of Western Australia can be compared to *Nepenthes*, as their ancestry, life history and presumably dispersal ability is similar. Most perennial and clonally reproducing Western Australian *Drosera* are self-incompatible and have higher seed abortion than self-compatible annual species (Stace et al. 1997). The mechanisms of self-incompatibility are diverse, suggesting multiple origins or continuing evolution and hence selection for outcrossing due to high genetic load in these poorly dispersing, clonal plants. Ancient hermaphroditic Nepenthaceae may have experienced similar pressure to reduce inbreeding.

Given the extreme nutrient and light limitation common in carnivorous plants (Givnish et al. 1984), sexual conflict may have been rather strong in the hermaphrodite ancestor, assuming that the costs of male and female reproduction differed (Barrett & Hough 2013; Obeso 2002; Zhang et al. 2014). Sexual selection in the form of male–male competition for ovules may favour higher flower numbers in males, whereas costs associated with the development of seeds under nutrient limitation might constrain females to a lower optimal flower number. While male and female reproductive costs and mate competition remain unexplored in carnivorous plants, in *Nepenthes* they may correspond to the difference in flower number between the sexes, which also varies between species (SI Table S1).

### Conclusions

The discovery of XY sex-determination in Nepenthaceae contributes to a better understanding of the diversity of plant sex determination systems and the molecular and ecological factors associated with dioecy and the evolution of sex chromosomes. As the foundation has been laid, future studies can address sexual conflict, X–Y chromosome divergence, and the identity of sex-determining genes. The species-rich radiation of *Nepenthes* lends itself to comparative studies. With the development of a simple molecular sexing assay, we also provide a tool for future studies on the ecological and physiological correlates of dioecy in *Nepenthes*, and anticipate that future work on *Nepenthes* will benefit from this resource. Our work exemplifies how sex-determination systems of non-model species can be studied and how statistically supported molecular markers suitable for the identification of sex can be developed without the need for prior genetic resources or existing breeding efforts.

## Acknowledgements

We thank N. Zemp for data of *Silene latifolia*. We are indebted to H. Luqman, C. Küffer, J. Mougal, K. Beaver, C. Morel, E. Nancy, A. Street, and T. Kropf for accessing *N. pervillei*. A.H.H. Tinggal and I. Daud with family are thanked for hospitality in Brunei, and L.W. Ngai for assistance with field work in Singapore. The Seychelles Bureau of Standards, the Seychelles Ministry of Environment and Energy, the Ministry of Industry and Primary Resources of Brunei Darussalam, the Agri-Food and Veterinary Authority and the National Parks Board of Singapore kindly granted permits for research, sampling and export of plant material. C. Michel is thanked for preparing sequencing libraries. S. Hartmeyer and U. Zimmermann kindly donated samples of sexed *Nepenthes* from their private collections. Sequencing and computation were carried out with the Genetic Diversity Center Zürich, the Quantitative Genomics Facility Basel, and the Functional Genomics Center Zürich. We thank T. Städler and two anonymous reviewers for valuable comments on earlier versions of the manuscript. This work was supported by ETH Zürich and partly supported through SNF grant no 31003A_160123. Keygene N.V. owns patents and patent applications protecting its Sequence Based Genotyping technologies.

## Author contributions

M.S. performed the research including data collection and analysis; T.U.G. and F.M. contributed logistic support and materials; M.S. and A.W. designed and interpreted the research and wrote the manuscript.

## Supporting Information

**Table S1** Literature survey of *Nepenthes* inflorescence dimorphism

**Table S2** Sex-associated markers

**Table S3** Overview table for sex-linked markers in *Nepenthes*

**Methods S1** Preliminary molecular sexing assay for *Nepenthes rafflesiana s.l*.

**Methods S2** Genotyping of *Silene latifolia*

**Methods S3** Male inflorescence transcriptome

**Methods S4** A molecular sexing assay for the genus *Nepenthes*

**Methods S5** Phylogenetic dating of *Nepenthes*

## Methods S1 Preliminary molecular sexing assay for *Nepenthes rafflesiana s.l*

An initial sequencing library contained a sufficient number of sexed individuals for *Nepenthes rafflesiana s.l*. and *N. gracilis* from Borneo. Collection and DNA extraction are detailed in the main text. We commissioned the Genomic Diversity Facility (Cornell University, Ithaca, NY, USA) with library construction and sequencing, following the GBS protocol (Elshire et al. 2011). After optimisation, the restriction enzyme Sfb1 was chosen and the library was sequenced for 100-bp single-end reads in two Illumina HiSeq lanes.

At this early stage, we employed a simpler version of the resampling approach to detect sex-specific loci, using the Stacks pipeline (Catchen et al. 2013) instead of the dDocent approach (Puritz et al. 2014) of genotyping. The populations module of Stacks was iterated over different combinations of real and permuted males and females, thereby revealing numbers and identities of likely sex-specific loci following the same logic as described in the main text (“privacy-rarefaction”). This attempt was successful in *N. rafflesiana* “typical form” (Borneo), but failed to identify any sex-specific loci in N. gracilis. We took the top 10 best candidate loci for *N. rafflesiana* “typical form” (Borneo) and designed PCR primers for validation of sex-specificity (PCR conditions as described in the main text). Two of these loci amplified from males exclusively (private gel band at expected size), as verified in all males that were used for the genotyping and several further samples that had not been used previously. The same markers also amplified specifically from known males but not females of *N. hemsleyana* and *N. rafflesiana* “giant form” (Borneo). However, these markers were unspecific for all other tested species (*N. ampullaria, N. bicalcarata, N. gracilis, N. mirabilis*). We consequently used these two markers to molecularly sex additional individuals of *N. rafflesiana* “typical form” (Borneo) and *N. hemsleyana* that were included in the later, full sample set genotyped with ddRAD-seq. To conclude, the sex of most *N. hemsleyana* and several of the *N. rafflesiana* “typical form” (Borneo) were determined not on the phenotype but molecularly with markers developed through this initial GBS dataset.

## Methods S2 Genotyping of *Silene latifolia*

Wild populations of *Silene latifolia* were sampled across Switzerland and leaves preserved by drying in silica gel. The phenotypic sex of individuals was recorded. After DNA extraction (Qiagen DNeasy Plant Mini Kit), a set of 95 samples was commissioned for library construction to the Genomic Diversity Facility (Cornell University, Ithaca, NY, USA), following the GBS protocol (Elshire et al. 2011). The restriction enzyme ApeKI was chosen after optimisation. This library was sequenced in two lanes of an Illumina HiSeq to ensure sufficient coverage.

The bioinformatics for *Silene* genotyping were identical to those employed for *Nepenthes*, as outlined in the main text respectively in *Scharmann et al* (*in revision*), i.e. *de novo* reference assembly following a modified dDocent pipeline (Puritz et al. 2014), read mapping, variant calling and quality filtering.

## Methods S3 Male inflorescence transcriptome

Plants of *Nepenthes khasiana* (*in vitro* propagated material from Borneo Exotics (Pvt) Ltd., Sri Lanka) were grown in a greenhouse where they flowered regularly. For the transcriptome of a male inflorescence of *N. khasiana* (length c. 2 cm, many buds of c. 1–3 mm diameter each), we extracted RNA using the Total RNA Mini Kit (Plant) (Geneaid Biotech Ltd, New Taipei City, Taiwan) with the “PRB” lysis buffer, which yielded undegraded high quality RNA (RIN 7.1; Plant RNA Nano Assay, Agilent Bioanalyzer). A cDNA library was generated (NEBNext Ultra Directional RNA Library Prep Kit for Illumina, New England Biolabs, Ipswich MA, USA) and sequenced in one lane of the Illumina MiSeq for 150 bp paired-end reads (GDC ETHZ). A total of 18.7 million PE reads were obtained, and a reference transcriptome was *de novo* assembled by the Trinity pipeline (Grabherr et al. 2011). Best ORFs were extracted by the Transdecoder script.

## Methods S4 A molecular sexing assay for the genus *Nepenthes*

Based on the evidence for male-specific genomic regions (non-recombing Y-chromosomeal region), we developed an assay to sex *Nepenthes* molecularly. Here we test it with phenotypically sexed individuals from 22 different *Nepenthes* spp. (Methods S4 Table S4–1). The assay is likely applicable to further *Nepenthes* spp., but we recommend to validate it using several phenotypically sexed individuals before application to a novel species.

**Methods S4 Table S4-1.** *Nepenthes* spp. with phenotypically verified sex used for broader taxonomic validation of a male-specific PCR marker

| species | N_male | N_female | source |
| --- | --- | --- | --- |
| *adnata* | 1 | 1 | cultivated |
| *albomarginata* | 1 | 3 | wild populations, cultivated |
| *ampullaria* | 2 | 2 | wild populations |
| *bicalcarata* | 2 | 2 | wild populations |
| *clipeata* | 1 | 0 | cultivated |
| *gracilis* | 3 | 3 | wild populations, cultivated |
| *hemsleyana* | 2 | 2 | wild populations |
| *khasiana* | 1 | 0 | cultivated |
| *maxima* | 2 | 0 | cultivated |
| *mira* | 0 | 1 | cultivated |
| *mirabilis* | 3 | 3 | wild populations |
| *mirabilis* var. *globosa* | 0 | 1 | cultivated |
| *pervillei* | 3 | 3 | wild populations |
| *petiolata* | 0 | 1 | cultivated |
| *rafflesiana* typical form Borneo | 3 | 3 | wild populations |
| *singalana* | 1 | 0 | cultivated |
| *talangensis* | 1 | 0 | cultivated |
| *tentaculata* | 1 | 0 | cultivated |
| *truncata* | 1 | 0 | cultivated |
| *veitchii* | 0 | 1 | cultivated |
| *ventricosa* | 0 | 1 | cultivated |
| *x trusmadiensis* | 1 | 0 | cultivated |

### DNA extraction

*Nepenthes* tissue contains substances that strongly inhibit PCR, as simple extraction protocols without purification steps (as suggested by (Hobza & Widmer 2008)) did not yield any amplification. We thus used the silica-column kit NucleoSpin Plant II from Macherey Nagel (Düren, Germany). For optimal yield, the tissue was completely powdered before the lysation step. To achieve this, tissue was flash-frozen in liquid nitrogen and then crushed in disposable, folded paper envelopes using pliers. The resulting coarse powder had to be still frozen, and was transferred to a 2 ml cryotube (Sarstedt No. 72.694.005, Screw Cap Micro Tube, 2 ml, PP, conical and skirted base) with three steel beads. On a shaker mill, cycles of shaking (up to 15 s) and flash-freezing in a liquid nitrogen bath were repeated until the material was a fine dust. Acceleration on the shaker mill was carefully adjusted to the maximum possible level that did not break the frozen cryotubes. Lysis buffer was added directly to the tissue dust without prior thawing. All other steps followed the kit instructions.

### PCR amplification of control and sex-specific sequences

The assay involves four primers: one pair targets a male-specific region (within the putative *Nepenthes* ortholog of the *Arabidopisis thaliana* DYT1 gene, 25100_L96_2_F: 5’-AATTCACTGATTCGGATCACG-3’; 25100_L96_294_R: 5’-CGATCGCGTCGCAAAGTATG-3’), while the other targets a sequence that is common to both sexes (the mitochondrial *cox1* gene; IP53: 5’-GGAGGAGTTGATTTAGC-3’; cox1.6KR: 5’-AAGGCTGGAGGGCTTTGTAC-3’; (Cho et al. 1998)). As suggested by (Hobza & Widmer 2008), the common target is used as an internal control of each reaction. It ensures that poor template quality or other technical issues are recognised as such, instead of confusion with true absence of the sex-specific target, i.e. the expectation for females. The two regions can also be amplified in separate reactions.

The reaction is performed in 15 *μ*l volumes containing 2.5 mM MgCl_2_, 250 *μ*M of each dNTP, 0.375 units of GoTaq DNA polymerase (Promega, Wisconsin, USA), 1x GoTaq Flexi buffer (Promega), 0.5 *μ*M of each of the four primers, and 1 *μ*l of template DNA extract (c. 5–20 ng/*μ*l). After initial denaturation for 2 min at 95°C, 30 cycles are run of denaturation for 30s at 95°C, annealing for 30s at 50°C and extension for 1 min at 72°C, followed by a final extension step of 5 min at 72°C (Thermocycler, e.g. Labcycler, SensoQuest, Göttingen, Germany).

### visualisation and scoring

PCR products are separated in 2% agarose gels and visualised by fluorescent dye (Methods S4 Fig. S4–1). Successful assays contain at least one strong band at 600–700 bp length, corresponding to the control *cox1* fragment. The presence of a strong band at c. 290 bp characterises male individuals, while females do not contain this band. Several other, weaker bands of different length may be present. These are likely unspecific products of the control primer pair, as we could never observe them when applying the sexing primer pair exclusively.

**Methods S4 Fig. S4–1.**
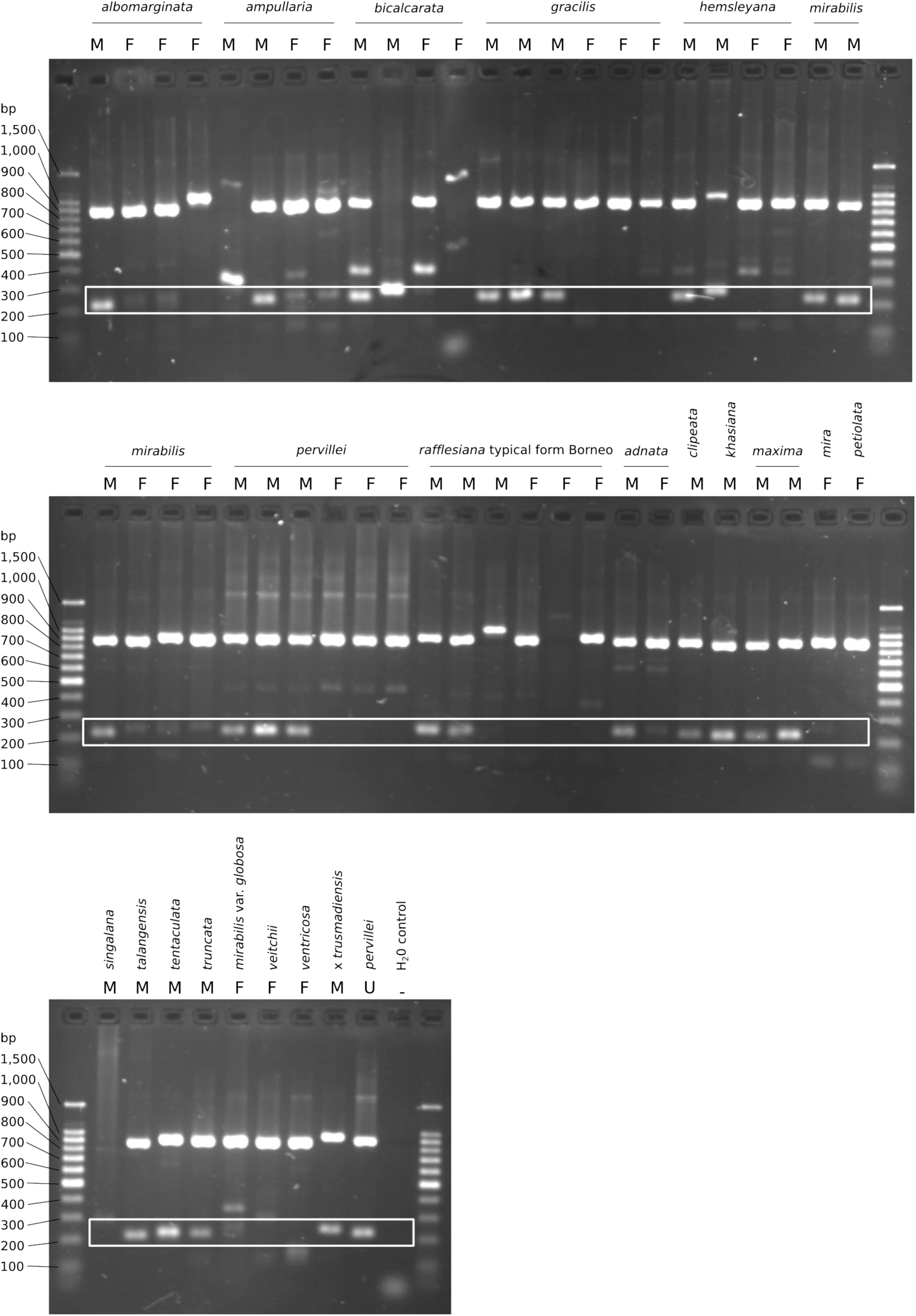
Electrophoretic gel confirming molecular sexing of *Nepenthes* spp. listed in Methods S4 Table S4–1. The region for male-specific bands is highlighted by the white box.

## Methods S5 Phylogenetic dating of *Nepenthes*

### Materials and method

We dated the genus *Nepenthes* by combining new transcriptome data for *Nepenthes* with previously published transcriptome data of the Venus Flytrap (*Dionaea muscipula*, (Bemm et al. 2016), a transcriptome-based phylogeny of Caryophyllales (Yang et al. 2015), and dates of Angiosperm diversification (Magallón et al. 2015). *Dionaea* represents Droseraceae, which is among the closest living sister lineages of *Nepenthes* (Brockington et al. 2009; Soltis et al. 2011).

In the first step, transcriptomes of 12 *Nepenthes* spp. (*Scharmann et al in preparation*) were assembled de-novo using Trinity (Grabherr et al. 2011). The raw assembly for *Dionaea* (v1.03) was downloaded from http://tbro.carnivorom.com/. We extracted candidate ORFs resp. peptide sequences with TransDecoder.LongOrfs v3.0.0 and TransDecoder.Predict. To reduce the sequence collections even further in a meaningful way, we retained only peptides that were similar (e-value <= 1e-5) to any gene from all available Eudicot plant genome assemblies (NCBI Genbank, accessed 6 June 2016).

In the second step, we emended a taxon-subset of the peptide sequence matrix for the 1,122 genes of (Yang et al. 2015) with orthologs from the *Dionaea* and *Nepenthes* transcriptomes. A custom python script was used to decompose the matrix by gene and taxon using the also available gene model file. The peptide sequences of “*Nepenthes alata* (WQUF)” were used to identify orthologs in the new transcriptomes by reciprocal best hit (blastp) with an e-value cutoff of 0.01. Third, a matrix was re-assembled with 21 of the original taxa and 13 newly added taxa, by globally re-aligning all ortholog peptide sequences of each gene using MUSCLE (Edgar 2004), and concatenation of the alignments. A new gene model file was generated in the process to allow partitioned analysis of the matrix. The new alignment was slightly longer than the original (550,076 instead of 504,850 sites), contained 34 taxa, and 21.7% gap characters. The 13 taxa we added showed very high sequence occupancy, each containing >1,000 of the original 1,122 genes of (Yang et al. 2015).

The maximum likelihood tree was reconstructed with the same method and partitioned by genes as before (RAxML -m PROTCATWAG -q; Yang et al. 2015). SH-like support was calculated using RAxML -f J option.

We then dated the divergence times on a pruned version of this tree (see below) using the RelTime algorithm (Tamura et al. 2012) as implemented in MEGA-CC (Kumar et al. 2012). RelTime is a non-Bayesian method for dating of phylogenentic trees that produces estimates similar to those from e.g. BEAST and MCMCtree, but it is orders of magnitudes faster and thus copes with genomics-scale alignments (Mello et al. 2017). The pruned tree contained only Brassicaceae (*Arabidopsis thaliana*) as the outgroup, and hence the alignment given to RelTime was also reduced with the same method as above (32 taxa, 550,360 sites, 22% gaps). We specified the WAG substitution model with 5 gamma-distributed rate categories and invariant sites. For 13 nodes that were also present in the Angiosperm time-tree of Magallón et al. (2015), we supplied absolute time calibrations in the form of upper and lower limits on age (Methods S5 Table S5–1).

Preliminary runs of RAxML and RelTime revealed that inclusion of Fabaceae, Rosaceae and Brassicaceae resulted in the same topology as retrieved by Yang et al. (2015), but enforcing these “Rosids” as a monophyletic outgroup caused RelTime to fit negative branch lengths near the root. However, reducing the outgroup to just Brassicaeae, RAxML found a rather different topology compared to Yang et al. (2015), and to this tree RelTime fitted negative branch lengths among the major lineages of Caryophyllales. Thus, to avoid biologically not interpretable negative branch lengths, we obtained a topology using the three outgroup taxa, but pruned this tree and the alignment to retain only Brassicaceae during RelTime dating.

### Results and discussion

We retrieved largely the same topology as Yang et al. (2015) for our subset of Caryophyllales taxa, with full SH-like LRT support for all nodes (tree not shown). The only exception, a grouping of Sarcobataceae as sister to Nyctaginaceae instead of Phytoloccaceae, occured in a lineage distant to *Nepenthes*. *Nepenthes* was monophyletic and grouped as sister to Droseraceae (*Dionaea muscipula*). This carnivorous lineage was sister to Frankeniaceae-Plumbaginaceae-Polygonaceae as reported before (Yang et al. 2015). The stem age of *Nepenthes* was estimated at 71.1 (CI 44.2 - 98.0) Mya, when it split from its presumed sister Droseraceae. The crown of *Nepenthes* is marked by the most basal species *N. pervillei*, and estimated here at 17.7 (CI 11.0 - 24.3) Mya.

However, we interpret these time estimates with great caution. First, the identity of the closest living relative of *Nepenthes* has not yet reached a consensus. Candidates are the lineage of sticky-leaf carnivores *Drosophyllum* and *Triphyophyllum* and several tropical lianas that appear to have lost carnivory secondarily (Heubl et al. 2006; Renner & Specht 2011), and the Droseraceae (Brockington et al. 2009; Soltis et al. 2011), or *Nepenthes* may even be basal to both of these (Brockington et al. 2015). We focussed on the Droseraceae because this was the only family for which transcriptome data was available.

Second, the divergence times that we took from the literature (secondary calibrations) may change in the future, as these were based on few genetic loci, fossils may be re-interpreted, and estimation methods change.

All previous attempts of molecular dating in *Nepenthes* (Merckx et al. 2015; Meimberg 2002) involved a presumed *Nepenthes* pollen fossil from the European Eocene (c. 50 million years ago; Krutzsch 1985). However, the attribution of this fossil to an ancestor of recent *Nepenthes* is not justified – it is larger than recent *Nepenthes* pollen but instead fits in the range of Droseraceae (Cheek & Jebb 2001). Thus, Krutzsch’s pollen fossils are at best indicative of the European Eocene presence of some lineage with Droseraceae-Nepenthaceae affinity but do not imply an age for modern *Nepenthes*.

**Methods S5 Table S5–1.** Absolute time calibrations as constraining upper and lower boundaries for the RelTime analysis, taken from Magallón et al. (2015). These are 95% confidence limits for the age (in million years) of the most recent common ancestor (MRCA) of 13 pairs of plant families studied by both Yang et al. (2015) and Magallón et al. (2015).

| MRCA of | min time | max time |
| --- | --- | --- |
| Cactaceae, Portulacaceae | 15.08 | 48.15 |
| Cactaceae, Talinaceae | 18.75 | 53.24 |
| Caryophyllaceae, Amaranthaceae | 50.34 | 88.56 |
| Cactaceae, Molluginaceae | 54.22 | 84.99 |
| Phytolaccaceae, Sarcobataceae | 63.24 | 73.41 |
| Plumbaginaceae, Polygonaceae | 65.63 | 78.21 |
| Aizoaceae, Nyctaginaceae | 72.5 | 77.87 |
| Phrymaceae, Solanaceae | 77 | 103 |
| Frankeniaceae, Plumbaginaceae | 83 | 101 |
| Caryophyllaceae, Physenaceae | 90 | 101 |
| Caryophyllaceae, Simmondsiaceae | 95.44 | 105.53 |
| Caryophyllaceae, Solanaceae | 114 | 123 |
| Brassicaceae, Nyctaginaceae | 120.87 | 126.49 |

